# Lf2 is a knotted homeobox regulator that modulates leaflet number in soybean

**DOI:** 10.1101/2025.06.20.660812

**Authors:** Chancelor B. Clark, Denise Caldwell, Qiang Zhu, Dominic Provancal, Austin C. Edwards, Qijian Song, Charles V. Quigley, Anjali S. Iyer-Pascuzzi, Jianxin Ma

**Affiliations:** Purdue University Department of Agronomy, West Lafayette, IN, USA; Purdue University Center for Plant Biology, West Lafayette, IN, USA; Purdue University Department of Botany and Plant Pathology, West Lafayette, IN, USA; USDA-ARS, Soybean Genomics & Improvement Laboratory, Beltsville, Maryland, USA

**Keywords:** allelic variation, gene cloning, genetic mapping, homeobox regulator gene, leaflet number

## Abstract

Variation in leaf complexity modulates light capture and is a target for crop enhancement. Soybean typically has compound leaves with three leaflets each, but a spontaneous mutation, designated *lf2,* possesses seven leaflets, offering a means to dissect the molecular mechanisms specifying leaflet number and assess its potential for soybean improvement. However, the developmental and genetic bases of the *lf2* mutation remain unknown. Here, we characterize the seven-leaflet phenotype and identify the mutation responsible for the phenotypic changes. Microscopic examination of leaf emergence sites revealed that the seven-leaflet phenotype arises in a two-step process: five leaflets form initially followed by secondary leaflet initiation at the margins of the central leaflet. Genetic mapping delineated *lf2* to a ∼2.5 Mb region at the start of chromosome 11. Fortuitously, integration of pedigree analysis with comparative analysis of genomic sequences from the region pinpointed a 2-bp deletion in the coding sequence of a gene, which is homologous to the Arabidopsis *KNAT7* encoding a *KNOTTED1-LIKE HOMEOBOX 2* transcription factor, as the sole candidate for *Lf2.* The deletion is predicted to result in disruption of the putative DNA-binding homeodomain. Expression of the wild-type allele of the candidate gene in the seven-leaflet *lf2* mutant restored the three-leaflet phenotype, while disruption of the wild-type allele through CRISPR-Cas9 editing induced extra leaflet formation. This study advances our understanding of leaflet formation in legumes and provides a template for utilizing compound leaf architecture to optimize photosynthetic efficiency and yield in soybean.

## Introduction

Leaves, the primary organs responsible for capturing light for photosynthesis, arise on the fringes of shoot apical meristems and exhibit a wide range of morphological differences within and among species (Bar and Ori, 2014). Leaf development can be classified into three stages: initiation, during which leaf primordia emerge from the periphery of the shoot apical meristem; primary morphogenesis, where the marginal leaf structures and lamina are formed; and differentiation, when the leaf tissues expand and mature to their final form (Bar and Ori, 2015). Compound leaves are those in which the lamina is subdivided into leaflets, in contrast to simple leaves in which the lamina is a single undivided unit (Bharathan and Sinha, 2001). These separate leaflets arise during primary leaf morphogenesis through the formation of distinct leaflet primordia (Mo et al., 2022). Compound leaves have evolved independently many times within the plant kingdom. They may hold ecological advantages for rapid growth of pioneering plant species, help plants tolerate herbivory, or be adaptations for response to seasonal drought, although evidence for each of these ecological functions is limited (Malhado et al., 2010). In plant species, differences in leaf and leaflet structure play a major role in determining light interception, the basis for photosynthesis and ultimately yield (Falster and Westoby, 2003). As a result, modifications to leaf structures are a major target of crop improvement as part of the fine-tuning of overall shoot architecture (Clark and Ma, 2023).

Compound leaf development results from the maintenance of the marginal blastozone (MB), a meristematic region capable of organogenesis (Blein et al., 2008). In many plant species, members of the *KNOTTED1-LIKE HOMEOBOX I (KNOXI)* gene family play a central role in maintaining the MB. In some legumes, particularly in the inverted repeat-lacking clade, this function is reported to be replaced in part by orthologs of *LEAFY* (*LFY*), which in *Arabidopsis* is associated with floral meristem identity. *KNOTTED1-LIKE HOMEOBOX 2 (KNOX2)* genes are also known to impact compound leaflet development, generally as negative regulators of leaf complexity which act antagonistically to *KNOX1* (Furumizu et al., 2015). Whereas *KNOXI* expression is closely associated with zones of undifferentiated cells, *KNOX2* genes are expressed in a broader array of differentiated tissues. In addition to compound leaf development, *KNOX2* genes have been implicated as important players in secondary cell wall development, stress response, and even nodule development (Di Giacomo et al., 2016; Ianelli et al., 2023; Wang et al., 2019).

Soybean, a critical crop for global oil and protein production, typically has two opposite simple (unifoliate) leaves at the first node after the cotyledons followed by single alternate compound leaves with three leaflets (trifoliate) at each subsequent node. Compound leaves are found in the majority of legume species, exhibiting diverse forms such as the soybean-like trifoliate leaflet arrangements in common bean (*Phaseolus vulgaris*), mung bean (*Vigna radiata*), and cowpea (*Vigna unguiculata*), the four opposite paripinnate leaflets in peanut (*Arachis hypogaea*), and the many alternate imparipinnate leaflets in chickpea (*Cicer arietinum*) (Liu et al., 2023).While the genetic basis of compound leaflet development is well characterized in some legumes, such as the model species *Medicago truncatula,* comparatively little is known about this trait in soybean.

Nevertheless, a recent study identified soybean *LFY* homologs, which, when knocked out, result in higher rates of simple leaves (Wang et al., 2025). Additionally, several studies have reported genetic loci resulting in multifoliate (i.e., more than the typical three) leaflets. *Lf1* is an incompletely dominant mutation that confers a high rate of pentafoliate (five leaflets) leaves at the normally trifoliate nodes (Fehr, 1972). It was mapped to a region with *Glyma.08g281900*, encoding an AP2-domain transcription factor, as the candidate gene (Jeong et al., 2017). There have been multiple other reports of loci independent of *lf1* associated with higher rates of multifoliate leaves (Wang et al., 2001; Orf et al., 2006; Wang et al., 2024). *lf2* is a recessive mutation that results in seven leaflets (heptafoliate) at the “trifoliate” nodes. *Lf1* and *lf2* display synergistic effects, with *Lf1Lf1;lf2lf2* individuals having 12-14 leaflets per “trifoliate” node (Fehr, 1972). Despite the *lf2* allele first being reported more than five decades ago, the genetic mutation or mutations responsible for the heptafoliate phenotype had not been identified. In this study we report the development pattern of the *lf2* heptafoliate phenotype, clone the gene underlying this locus, and identify potential downstream targets of Lf2.

## Results

### Characterization of the *lf2* Phenotype

To understand the *lf2* seven-leaflet soybean trait, we evaluated the lines known to possess it. The seven-leaflet trait was originally reported as a spontaneous mutation (i.e., the sudden appearance of an individual plant with seven leaflets) in the trifoliate cultivar Hawkeye (Fehr 1972), and this line was designated as T255(PI 548232). This trait was then backcrossed into the cultivar Clark (PI 548533),to develop the *lf2/lf2* Clark near isogenic line (NIL) L73-1087 (PI 547580) one of many resources developed as part of the USDA soybean isoline collection (Gilbert et al., 2023). L73-1087 has three leaflets on each of the opposite leaves at the first node, in contrast to the wild-type unifoliate single leaflet, and seven leaflets on each subsequent node in contrast to the wild-type trifoliate (Fig. 1A; Supplementary Fig.S1). The phenotype is consistent, although occasionally 6, 8, or 9 leaflets were observed instead of seven on individual nodes. Unlike in other multifoliate soybean mutants, trifoliate leaves are never observed in L73-1087 after the first node. In general, the total leaf area is greater in L73-1087 than in typical wild-type trifoliate soybeans.

**Figure 1.**
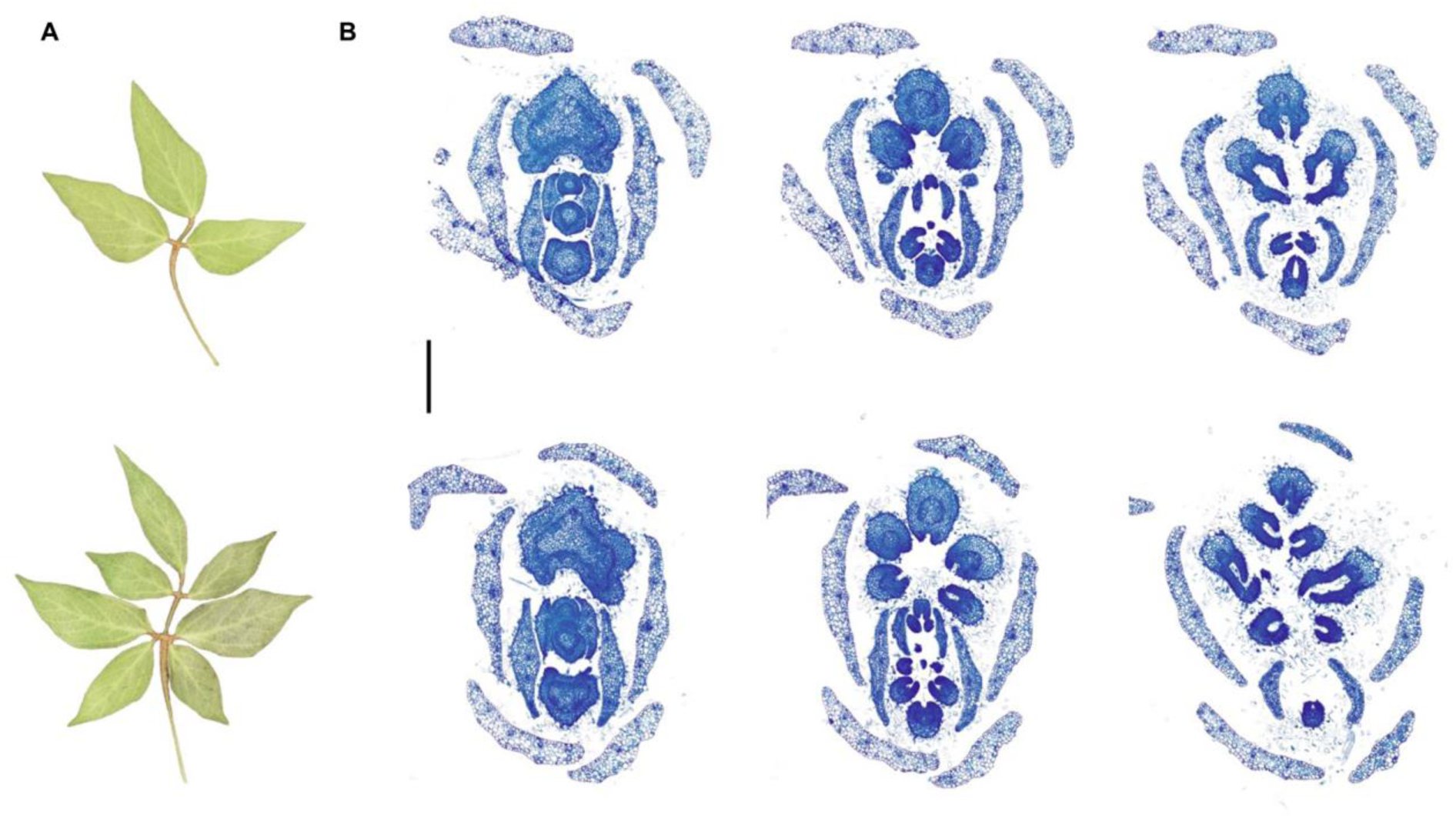
Characterization of *lf2* Mutant. **A)** Mature leaves of Williams 82 (Lf2;Lf2, top, trifoliate) and L73-1087 (lf2;lf2, bottom, heptafoliate). **B)** Light microscope images of developing leaf primordia. Williams 82 (top) progresses from a single leaf primordium to a compound trifoliate leaf. L73-1087 (bottom) progresses from a single leaf primordium to a palmately compound leaf with five leaflets. The central leaflet then undergoes secondary leaflet formation at the margins for a total of seven leaflets. The scale bar is 40 μm.

To trace the development of the seven-leaflet phenotype, we performed light microscopy of cross sections of the meristem (Fig. 1B). In trifoliate soybean leaflets, the terminal and two lateral leaflets emerge nearly simultaneously from the leaf primordium. In the heptafoliate mutant, initially five leaflets emerge in a palmately-compound fashion, corresponding to the two lateral leaflets on each side and one central leaflet attached to the rachis. Subsequently, the central leaflet undergoes secondary leaflet formation as two lateral leaflets emerge from the leaflet margins. This secondary leaflet formation at the central leaflet follows a reiterative pattern that mimics the normal formation of trifoliate leaves in wild-type soybean.

### Genetic Inheritance Pattern of *lf2*

To understand the inheritance pattern of the *lf2* mutation, we developed a mapping population by crossing L73-1087 to Williams 82 (Lf2/Lf2, trifoliate). The F_1_ hybrids had wild-type unifoliate leaves at the first node and most subsequent nodes had wild-type trifoliate leaflets but each F_1_ plant had an occasional node with four or five leaflets. In the F_2_ population, 89 individuals had seven leaflets per leaf and 328 individuals had three or mostly three leaflets, consistent with a single gene modulating leaflet number in this population (χ^2^ for 3:1 = 2.9744, p-value = 0.084594). Trifoliate is incompletely dominant in this population, as evidenced by the occasional appearance of leaves with four, five, or six leaflets or extra lobes in the F_1_ and heterozygous F_2_ individuals.

### The *Lf2* Locus is on the long arm of chromosome 11

In order to detect the genomic location of the gene responsible for the *lf2* 7-leaflet phenotype, we carried out bulked segregant analysis (BSA) of typical trifoliate or heptafoliate individuals in the F_2_ population. BSA places the *Lf2* locus at the beginning of chromosome 11 (Fig. 2 A). Markers ranging from 2119662 base pairs (bp) to 2574195 bp were homozygous for the L73-1087 genotype in the heptafoliate bulked sample (Supplementary Table S1). However, there were no markers displaying polymorphism between Williams 82 and L73-1087 upstream of bp 2111286, so we must consider the mapped region to be from 0 to 2574195 bp on chromosome 11 based on the principle of BSA. No other regions were homozygous in either bulked sample. *Lf2’s* placement on chromosome 11 is consistent with older studies showing it belongs to classical linkage group 16 and molecular linkage group B1 (Devine, 2003; Seversike et al. 2008).

**Figure 2.**
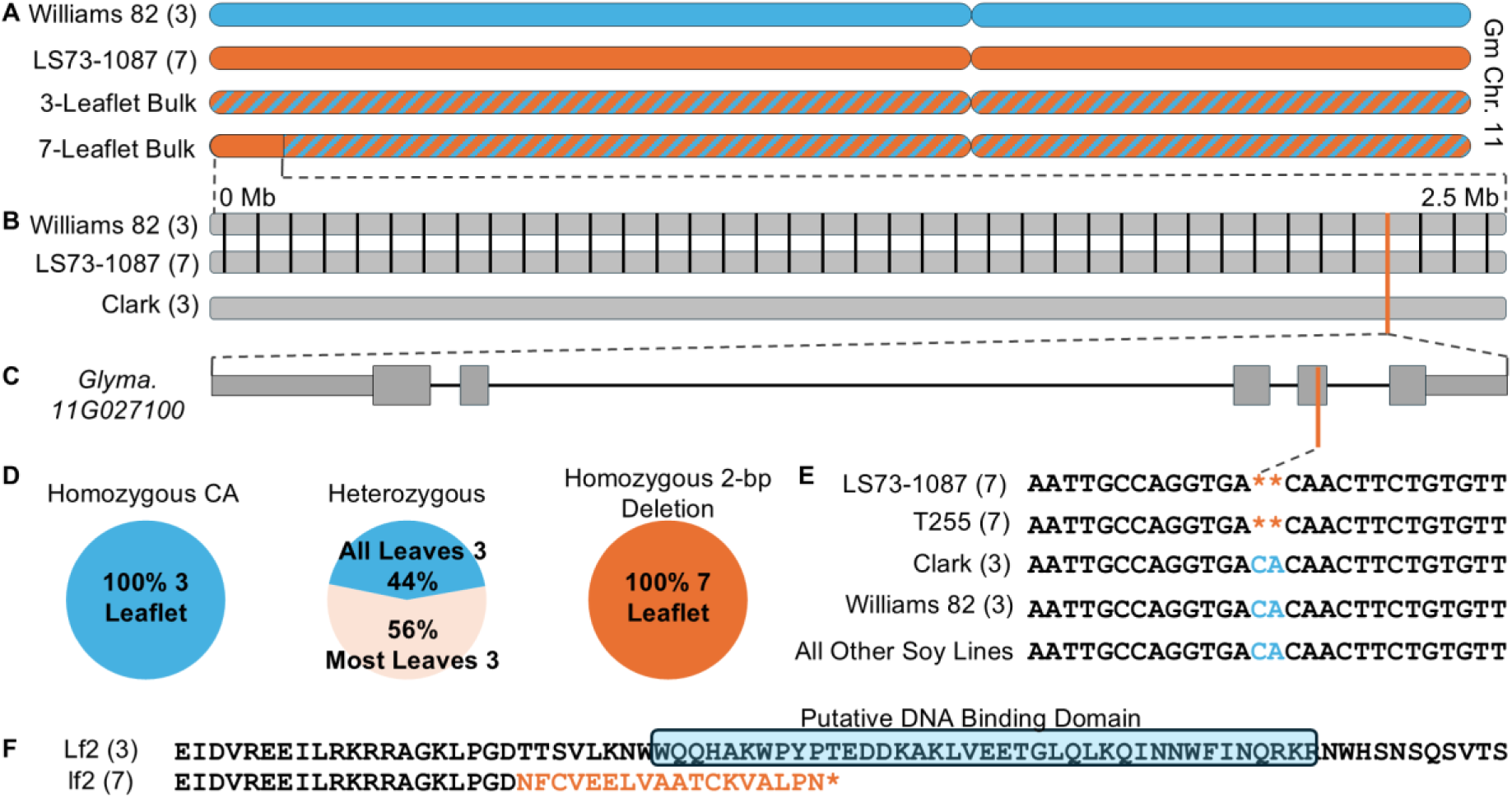
Identification of the Genetic Basis for *lf2.* **A)** The location of the lf2 region on chromosome 11 as defined by BSA. The blue chromosome segments correspond to Williams 82, the orange to L73-1087, and the blue-orange stripes show heterozygous. **B)** Genetic variation between Williams 82, L73-1087, and Clark in the lf2 region defined by BSA. Each black bar represents a polymorphism between Williams 82 and L73-1087 while the orange bar shows the only polymorphism between L73-1087 and Clark. **C)** The gene structure of *Glyma.11G027100.* The orange bar shows the location of the 2-bp deletion in the fourth exon. **D)** Pie charts showing the phenotypic distribution for each allele at the 2-bp deletion site demonstrating perfect correspondence between the genotype and phenotype **E)** Sequence alignment at the site of the 2-bp deletion, the only mutation that can distinguish the 2 *lf2;lf2* 7-leaflet lines from their *Lf2;Lf2* trifoliate counterparts. **F)** The consequence of the 2-bp deletion on the predicted protein sequence, eliminating the putative homeodomain DNA Binding Domain. The putative homeodomain is highlighted in light blue while the changed amino acids in the mutant are shown in orange.

### A 2-bp Deletion in a Knotted Homeobox Transcription Factor is the Likely Causal Mutation for *lf2*

Since L73-1087 is a near isogenic line developed by backcrossing the seven-leaflet trait from a spontaneous 7-leaflet mutant line observed in the cultivar Hawkeye (T255), into the recurrent trifoliate parent Clark, we thought we would be able to observe this introgression in the region defined by BSA on chromosome 11. Surprisingly, publicly available SNP genotyping data for L73-1087, Clark, and T255 show introgressions of T255 alleles on chromosomes 2, 3, 15, and 17, but not on chromosome 11 (Supplementary Table S2). This can be explained by the half-sibling relationship between Hawkeye (Mukden × Richland) and Clark (Lincoln × Richland), which could both have inherited the region containing *Lf2* from Richland, hiding the introgression (Supplementary Figure S2). As a result of the low amount of genetic variation between Clark, Hawkeye, and Williams 82 in the region defined by BSA, we suspected that we could easily find only one or a small number of mutations which met the criteria of occurring in the heptafoliate lines L73-1087 and T255 but not in the trifoliate lines from which they were derived, Clark and Hawkeye, nor in Williams 82.

To find a mutation that could distinguish the three and seven leaflet lines, we carried out whole genome resequencing of L73-1087. In the region defined by BSA, only a single mutation could distinguish the heptafoliate L73-1087 from Clark, Hawkeye, and Williams 82, a 2-bp deletion in the fourth exon of *Glyma.11g027100* (Fig. 2 B; Fig. 2 C). While there are 38 polymorphisms between Williams 82 and LS73-1087, only the 2-bp deletion distinguishes LS73-1087 from Clark (Fig. 2 B). This deletion is predicted to cause a frameshift for eighteen amino acids and then a premature stop codon, shortening the annotated protein from 279 to 232 amino acids. *Glyma.11g027100* encodes a Class II KNOTTED-like homeobox (KNOX II) transcription factor, members of which are associated with leaf and leaflet development in many plant species (Wang et al. 2023). The frameshift and truncation disrupt the putative DNA-binding homeodomain ranging from amino acids 223 – 261 (Fig. 2 F). The loss of the homeodomain consisting of three alpha helices can be easily visualized in the predicted protein structures of the mutant and wild-type generated by AlphaFold (Supplemental Fig.S3).

To confirm the association between the 2-bp deletion and the phenotype, we carried out Sanger sequencing of T255 and our F_2_ mapping population. As expected, the mutation is also present in T255, the original line in which the seven leaflet phenotype was reported. (Fig. 2 E). In the F_2_ mapping population, there was a perfect correspondence between the mutation and the leaflet phenotype, with all lines possessing the deletion having seven leaflets and all lines without the deletion having three leaflets (Fig. 2 D). Multifoliate (4 or more) leaflets were observed on at least one node in about half (56%) of heterozygous lines, consistent with a single gene model where trifoliate is incompletely dominant over heptafoliate.

Since the 7-leaflet trait originated as a spontaneous mutation in the cultivar Harosoy, and because the 7-leaflet phenotype is not described in lines besides T255 and L73-1087, we should not expect to find the 2-bp deletion or other mutations in *Glyma.11g027100* in publicly available soybean resequencing data. To confirm this, we checked ∼3000 resequenced wild and domesticated soybean accessions. We find a complete absence of mutations that change the amino acid sequence of this gene in any line. This further provides evidence of the importance of *Lf2* in regulating normal leaflet number in soybean.

### Validation of *Lf2*

To validate *Glyma.11g027100* as *Lf2,* the CDS of the Williams 82 *Lf2* allele was transformed into the heptafoliate line L73-1087 under the influence of the 35S promoter. The Williams 82 *Lf2* allele was able to rescue the leaflet number phenotype (Fig. 3A). Two independent lines had all or nearly all (one leaf with four leaflets were observed on one line) while a third event which was genotyped as positive but in which low expression of the construct was detected showed no difference from LS73-1087 (Fig. 3A,B,D).While the complementation nearly eliminated the multifoliate phenotype, we did not observe further reductions in leaf complexity (e.g., simple leaves or leaves with less than three leaflets), clarifying the specific role of *Lf2* in maintaining leaflet number.

**Figure 3.**
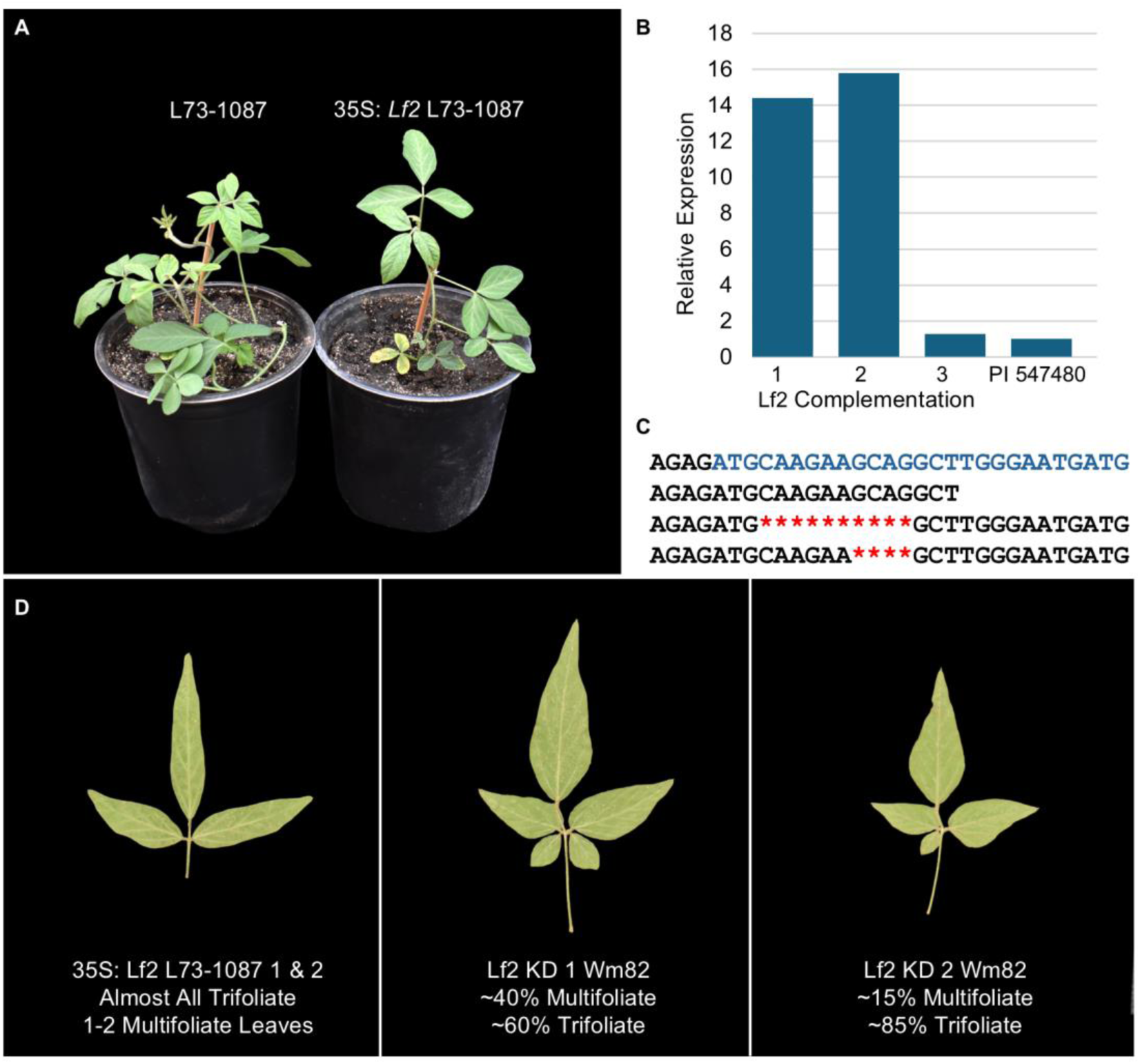
Validation of *Glyma.11g027100* as *Lf2*. **A)** Overexpressing *Glyma.11G027100* CDS in L73-1087 restored the trifoliate phenptype. **B)** Expression of *Glyma.11G027100* in three complementation lines. The gene was highly expressed in lines 1 and 2, leading to nearly complete complementation, but line 3 displayed expression not significantly different than the control, suggesting it had zero or very low expression and consequently no change in phenotype. **C)** Alignment of the knockdown line edited sites (bottom two rows) with the beginning of the annotated CDS of *Glyma.11g027100* (top row, with the CDS shown in blue) and the guide RNA (second row). **D)** Summary of leaflet phenotypes in complementation and knockdown lines with representative example leaflets

CRISPR-Cas9-mediated knockdown was used to further validate *Glyma.11g027100* as *Lf2* (Fig. 3C). Two independent lines with deletions near the annotated start codon were generated. These lines display high rates of four and five leaflets, as well as other leaf abnormalities such as extra lobes (Fig. 3D; Supplemental Fig. S4). One event with a ten bp deletion displayed 40% of leaves with four or five leaflets while a separate event with a four bp deletion displayed 15% of leaves with four leaflets. RNA sequencing data shows that *Glyma.11g027100* has alternative transcripts which were unaffected by deletions at this site, allowing the gene function to be partially maintained (Supplemental Fig. S5). Aside from the changes to leaflet number, no other obvious phenotypic changes were observed in the complementation and knockdown lines. The complementation and knockdown results, combined with the genetic mapping data provide compelling evidence that *Glyma.11g027100* is *Lf2*.

### DAP-Seq Reveals Potential Downstream Targets of *Lf2*

To confirm the subcellular localization of *Glyma.11g027100*, we amplified the CDS and added a GFP tag for expression in tobacco. GFP signal was observed exclusively in the nucleus, consistent with its expected role as a transcription factor (Fig. 4A). Having confirmed the nuclear localization of *Lf2,* we performed DNA Affinity Purification Sequencing (DAP-Seq) to identify potential targets of Lf2 regulation. A total of 344 binding peaks were identified in promoters or gene bodies, including several with known or suspected functions in the specification of compound leaf development. Analysis of the overrepresented motifs across two replicates found that TGACAKBT was the most common shared sequence (Fig. 4B), found in more than 60% of significant binding peaks and consistent with homeobox transcription factor binding sites identified in other plant species (Ou et al., 2022). Notably, a binding site was found in the promoter region of *Glyma.08G281900,* previously identified as the candidate to be the leaflet number regulator *Lf1.* The peak of the binding site in the *Glyma.08G281900* promoter contains the motif TGACAGCT (Supplemental Fig.S6). Another gene which could be bound by *Lf2* potentially associated with compound leaf development is *Glyma.04G128800,* a homolog of the *A. thaliana* C2H2 zinc-finger *SERRATE (SE)* (Prigge and Wagner, 2001). Several genes related to auxin, a key player in leaflet development, were also found to be bound by *Lf2,* including *Glyma.14G120900* and *Glyma.19G161000,* homologs of the *A. thaliana* auxin efflux transporter *PIN6* and the AUX/IAA transcriptional regulator *ATAUX2-11*, respectively. The complete list of genes associated with binding peaks can be found in Supplementary Table S3.

**Figure 4.**
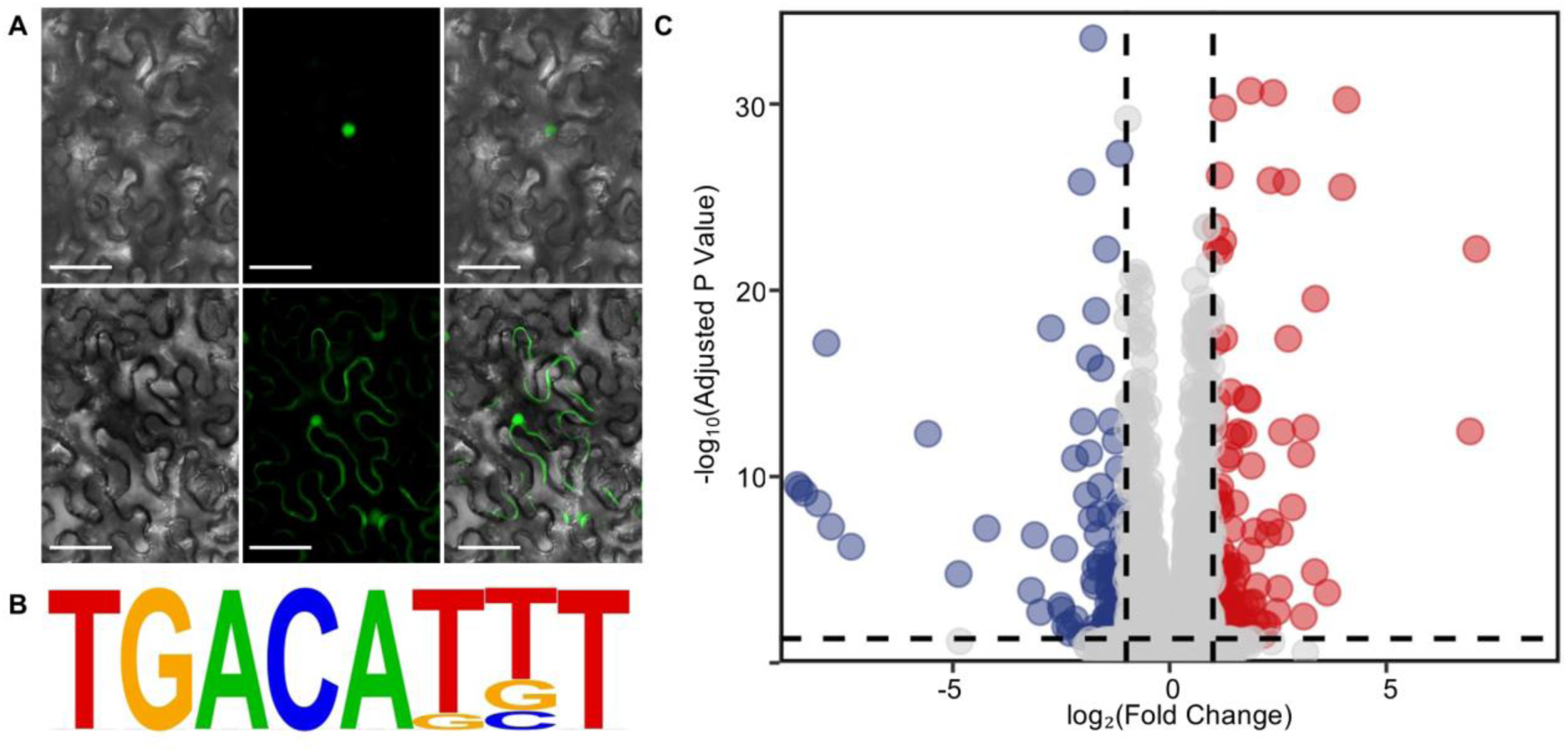
Regulatory profiling of *Lf2.* **A)** Nuclear localization of *Lf2* in *N. tabacum*. Top from left to right shows Lf2 Brightfield, GFP, and Merged. Bottom row from left to right shows control GFP brightfield, GFP, and Merged. **B)** The sequence logo corresponding to the putative binding site of *Lf2*, found in more than 60% of DAP-Seq Peaks. **C)** Volcano plot of total RNA sequencing results from Clark (*Lf2Lf2*) and LS73-1087 (*lf2lf2*). Genes significantly upregulated in the lf2 mutant are shown in red while genes significantly downregulated are shown in blue.

### RNA Sequencing of Clark *Lf2/lf2* Near Isogenic Lines Sheds Light on Downstream Effects of *Lf2*

To gain further insight into how the loss of function *lf2* allele changes patterns of gene expression, we performed total RNA sequencing of V1 stem tips of the Clark (*Lf2;Lf2*) and L73-1087 (*lf2;lf2)* near isogenic lines. A total of 246 genes were found to be differentially regulated, of which 130 were significantly upregulated and 116 significantly downregulated in L73-1087 (Fig. 4C). Among these were homologs of many genes known to be associated with compound leaflet development in other plant species. One gene significantly upregulated in the heptafoliate line is *Glyma.01G198700,* the most similar soybean gene to *M. truncatula PALM1,* which encodes a C2HC zinc fingers superfamily protein well characterized for its essential role in compound leaf development (Chen et al., 2010; Ge et al., 2010). An *A. thaliana KNAT6* homolog, *Glyma.18G141200*, was downregulated in the heptafoliate line. Overexpressing the KNAT6 homolog *MtKNOX2* in *M. truncatula* results in increased leaflet number (Lu et al., 2024). Three homologs of *A. thaliana TCP1*, part of a key transcription factor family for compound leaf development in many species including tomato (Bi et al., 2024), were also found to be upregulated in the heptafoliate line. Finally, two *BEL1* homeodomain genes, *Glyma.02G060100* and *Glyma.05G210300* were found to be differentially regulated. A BEL1 homolog in *M. truncatula*, MtPINNA1 (He et al., 2020), is a major player in maintaining trifoliate leaf development and BEL-like, KNOX, and TCP transcription factors are reported to work together to regulate leaf margin development in *A. thaliana (*Yu et al., 2021).

*Glyma.11G027100*, along with each of its three duplicate paralogs, was also found to be significantly upregulated, although the log_2_(fold change) was only about 0.7 for each. This may represent compensatory upregulation in response to the loss of function. There was not a strong correspondence between the genes identified to be bound by DAP-seq and those differentially expressed in the RNA-seq data, although in each case genes involved in similar pathways were identified. This is consistent with our RNA-seq data - with samples taken from whole stem tips - reflecting downstream transcriptional effects of *lf2* loss of function rather than direct targets. The complete list of differentially expressed genes is given in Supplementary Table S4.

## Discussion

While not as ubiquitous among compound leaf development mutants as KNOX 1 homeoboxes, which are positive regulators of leaf complexity across the plant kingdom, KNOX 2 homeoboxes have been implicated in modulating this trait in several species. In fact, knocking out the putative ortholog of *Lf2* in the model legume species *Medicago truncatula, MtKNOX4,* also increased leaflet number, from the typical three to five in the mutant (Wang et al., 2023). This phenotype is similar to that caused by *lf2,* but distinct in at least three ways. Firstly, essentially every leaf in the *lf2lf2* mutant lines has increased leaflet numbers while only around half of the *Mtknox4* leaves displayed increased leaflet numbers. Secondly, *Lf2* changes the leaves at the first node from simple to compound, while *Medicago truncatula* has trifoliate leaves at the first node. Finally, *Lf2* helps to regulate both primary and secondary leaflet formation in soybean, leading to seven leaflets rather than the five observed in the *Mtknox4* loss-of-function mutants. This shows at least a partial divergence in gene function between soybean and *M. truncatula* and suggests that further study of compound leaf development in soybean is warranted.

Our DAP-Seq and RNA-Seq results provide a starting point for understanding how *Lf2* maintains trifoliate leaflets in soybean. The differences between the genes identified by DAP-seq and those differentially expressed between the near isogenic lines likely stem from the tissues which were collected. Based on our light microscope observations of the developing leaflet primordia and what is known about leaflet development in other plant species, the specification of leaflet number occurs early in leaflet development. Furthermore, the region critical for leaflet organogenesis, the marginal blastozone, only makes up a fraction of the developing leaflet. Therefore, compared to the total amount of tissue collected in the stem tips, the likely site of *Lf2* function in regulating leaflet number is quite small, and the ability of traditional RNA seq to capture the direct transcriptional regulation of *Lf2* may be limited. Future work to help resolve this using single cell RNA sequencing or other similar techniques may be valuable. Nonetheless, the appearance of numerous homologs of genes known to be involved in compound leaf patterning across the plant kingdom suggests these samples were beneficial in capturing the indirect downstream effects of the *lf2* loss-of-function allele.

Understanding the genetic basis for differences in leaflet number could prove useful for soybean improvement. A study of *Lf2/lf2* near isogenic lines showed that the 7-leaflet trait had no effect or a negative effect on yield at higher planting densities (Seversike et al., 2009). However, this study also found that at low densities the 7-leaflet trait could have advantages for yield due to increased light interception. This is consistent with 7-leaflets being beneficial when light interception is a limiting factor and being a waste of resources once canopy closure is achieved. Rapid canopy closure and efficient light interception drive yield in many crop species (Sreekanta et al., 2024). Increased leaflet number on upper leaves likely contributes to increased shading of lower leaves, cancelling out potential benefits from greater light interception. With the rapid development of gene editing technology such as CRISPR/Cas9 and methods for precise tracking of the spatiotemporal expression of genes, we are less constrained to only selecting between naturally occurring leaflet number phenotypes. We speculate that a putative soybean leaflet number ideotype of heptafoliate lower leaves and trifoliate upper leaves could strike the balance of increased light interception early in the growing season before canopy closure without causing increased shading or wasting resources in the upper canopy. This phenotype could be achieved through growth-stage-specific downregulation of *Lf2* or other genes in the leaflet development pathway. Having increased leaflet number only at the lower nodes of the plant would be analogous to reduced upper leaflet angle in maize, which facilitates the efficient capture of light at high planting densities (Mantilla-Perez and Fernandez, 2017). Manipulation of soybean leaflet number, perhaps in combination with other soybean shoot architecture traits like branch angle (Clark et al., 2022; Virdi et al., 2023), could be a path to optimized soybean shoot architecture. Much remains unknown about compound leaflet development in soybean, providing many opportunities for deeper dissection of this trait.

## Materials and Methods

### Plant Materials and Mapping Population Development

The parent lines used for mapping were Williams 82, a trifoliate cultivar which was used for the original soybean reference genome (Schmutz et al. 2010) and L73-1087 (PI 547580), a heptafoliate near-isogenic line which was developed by backcrossing the 7-leaflet trait from the line T255 (PI 548232) into the cultivar Clark (PI 548533), one of many resources developed as part of the USDA soybean isoline collection (Gilbert et al., 2023). T255 in turn was reported as a spontaneous mutation (i.e., the sudden appearance of an individual plant with seven leaflets) in the trifoliate cultivar Hawkeye (Fehr 1972).

The mapping population was developed by crossing Williams 82 to L73-1087, using L73-1087 as the pollen parent and Williams 82 as the female parent. 15 F_1_ hybrid lines were confirmed by the appearance of purple flowers and black pods, dominant traits possessed by L73-1087 but not by Williams 82. A total of 417 F_2_ individuals were grown in the field at the Purdue Agronomy Center for Research and Education (ACRE), in West Lafayette, Indiana, United States of America.

### Histology and Light Microscopy

Soybean meristems (2–5 mm) were excised from the region where the first true leaves emerged. The stem was cut just below the leaf emergence site and approximately 3 mm above it to isolate this zone. Larger leaves were removed by severing them at the petiole.

Samples were immediately fixed in a solution of 1.5% glutaraldehyde and 2% paraformaldehyde in 0.1 M sodium cacodylate buffer (pH 6.8) under low vacuum for 2 h, as previously described (Caldwell et al., 2017; Caldwell & Iyer-Pascuzzi, 2019). Samples were washed in a buffer under vacuum and then subjected to a graded dehydration series using ethanol and tert-butyl alcohol (TBA), with intermediate low-vacuum treatments. Following three changes in 100% TBA over 24 h, tissues were infiltrated with molten paraffin at 54°C. Samples were embedded in molds, oriented perpendicularly, and solidified at room temperature.

Paraffin blocks were mounted to embedding rings, trimmed, and sectioned at 12 µm thickness using a rotary microtome. Sections were floated on the water at 34°C, mounted to slides, dried, deparaffinized, and stained with 0.05% toluidine blue using a graded xylene–ethanol series. Stained samples were mounted with Permount and coverglass and then imaged on an Olympus BX43 compound microscope equipped with a DP80 Dual CCD camera.

### Phenotyping

Leaflet numbers were first phenotyped at the V2 stage when the first set of opposite “unifoliate” leaves and two sets of “trifoliate” leaves could be observed. These initial phenotypes were confirmed at the V5 stage.

### DNA Isolation and Bulked Segregant Analysis

F_2_ leaf tissue was collected at the V5 stage and DNA was isolated in 96-well plates using a modified CTAB method (Mace et al., 2013). For bulked segregant analysis (BSA), leaves from fifteen plants with seven leaflets and fifteen plants with three leaflets were bulked together and ground to a fine powder using liquid nitrogen. The ground bulked samples, as well as samples of Willams 82, L73-1087, and Clark, were then isolated using the same CTAB method described above.

The five samples were then genotyped with the Illumina Infinium SoySNP50K BeadChip (Song et al., 2013). Each SNP genotype was classified manually using the GenomeStudio software from Illumina. The BSA mapping region was defined as the area in which the recessive (7 leaflet) bulk was homozygous for the recessive (L73-1087) parent genotype, according to the principle of BSA.

### Resequencing of L73-1087

The same L73-1087 DNA sample used for SNP genotyping was also used for resequencing. Sequencing was carried out by the Purdue Genomics Core (West Lafayette, IN, USA) and reads were sequenced to a depth of 20x coverage. Reads were aligned to the Williams 82 second annotation reference genome using the Burrows-Wheeler Alignment tool (BWA) (Li and Durbin 2009) and the data was processed using the SAMtools suite (Li et al., 2009). The region on chromosome 11 defined by BSA was compared to publicly available sequence data for Clark and Williams 82 manually using the Integrated Genome Viewer (IGV). Hawkeye’s genotype was inferred using publicly available resequencing data for its parents, Richland and Mukden.

### Subcellular Localization of *Glyma.11G027100*

Total RNA from Williams 82 young leaves was reversed transcribed using MMLV Reverse Transcriptase (Promega, Madison, WI, USA). *Glyma.11G027100* CDS was amplified with Q5 High-Fidelity Polymerase (New England Biolabs, Ipswitch, MA, USA). The CDS was then inserted into the vector pFM3100, under the influence of the Cauliflower mosaic virus 35S promoter and containing an N terminal green fluorescent protein (GFP) tag. This construct was transformed into the *Agrobacterium tumefaciens* strain EHA-105. The Agrobacterium culture was diluted in magnesium chloride and injected into *Nicotiana tabacum* leaves. Two days after inoculation, transformed leaves were observed using an Olympus BX43 light microscope (Olympus Corporation, Hachioji, Tokyo, Japan) illuminated with an X-Cite SERIES 120 Q Illumination System (Excelitas, Pittsburgh, PA, USA).

### Sanger Sequencing of Causal Mutation

The following primers flanking the 2-bp deletion were used to check the genotypes of T255 and the F_2_ mapping population. Forward: TCAAAACTTCACGTGCTAAACTC. Reverse: ATCTCTGCAGGGTTTCAAGTCA. PCR products were sent to Eurofins Genomics Sequencing Laboratory (Louisville, KY) for sanger sequencing using the forward primer listed above.

### CRISPR Cas9 Mediated Knockdown of *Lf2*

Guide RNA target sequence AGAGATGCAAGAAGCAGGCT was ligated into the CRISPR/Cas9 Construct pGES201 (Bai et al., 2020). Successfully edited plants were identified by sanger sequencing of the target sites at Eurofins Genomics (Louisville, KY). The following primer pair was used for amplification of the target site, Forward: TTTCTGACCTCGCTTCCATCC; Reverse: GTTGTCAAGTTCTTGGCGGT.

### *lf2* complementation

Total RNA from Williams 82 young leaves was reverse transcribed using MMLV Reverse Transcriptase (Promega, Madison, WI, USA). *Glyma.11G027100* CDS was amplified with Q5 High-Fidelity Polymerase (New England Biolabs, Ipswitch, MA, USA) and inserted into the vector pPTN1171 (Ping et al. 2014) using the ClonExpress II One Step Cloning Kit (Vazyme, Nanjing, China). The following nested primers were used for amplification and insertion: outer F: ACCCTATATTATTCAACCCAATCCT, outer R: TGCTTGCTAGCTGTTCCACA, inner F: TTTACGAACGATAGCATGCAAGAAGCAGGCTTGGGA, inner R: TGATTTTTGCGGACTCTACCTCTTGCGTTTGGACT.

### Soybean transformation

The complementation construct was transformed into L73-1087, and the CRISPR/Cas9 knockdown construct was transformed into Williams 82 via the *Agrobacterium tumefaciens* mediated cotyledonary node method and regenerated through sterile soybean tissue culture as described in Wang et al., 2024.

### DNA affinity purification sequencing (DAP-seq)

Williams 82 genomic DNA (isolated as described above) and the pPTN1171 plasmid containing *Lf2* CDS were sent to CD-Genomics (Shirley, NY, USA) who carried out DAP-Seq Genomic Library Preparation, DAP-Seq Protein Expression, and DAP seq binding assay and sequencing. Reads were aligned to the Williams 82 reference genome, second annotation first version, using BWA-MEM (Li 2013). Aligned reads were sorted, indexed, and filtered for uniqueness using SAMtools (Li et al. 2009). 2 technical replicates were compared to the input genomic DNA and DAP-Seq Binding Peaks (*q* value < 0.05 and fold enrichment > 2) were called using MACS2 (Feng et al. 2012). HOMER (Heinz et al. 2010) was used to find the most overrepresented sequences within the DAP-Seq Peaks to identify potential binding motifs.

### RNA Sequencing

RNA from 10 V0 Clark and L73-1087 stem tips for each of three replicates was isolated with Trizol Reagent (Invitrogen, Carlsbad, CA, USA). RNA quality and concentration was checked with Nano Drop (Thermo Fisher Scientific, Waltham, MA, USA ). DNA was removed using the Turbo DNA Free Kit (Thermo Fisher Scientific, Waltham, MA, USA). Samples were sent to BMK Gene (Beijing, China) for total RNA sequencing.

Uniquely mapped reads were aligned to the Williams 82 (second annotation first version) reference genome using HISAT2 (Kim et al., 2019). Reads were formatted using SAMtools. Read counts were obtained using subread featureCounts (Liao et al., 2014). Differentially expressed genes were identified using DEseq2 (Love et al., 2014) and filtered to include only those with base mean above 10, adjusted *p*-value (FDR) below 0.05 and an absolute log₂ fold change ≥ 1.

## Data Availability

The transcriptomic, genomic, and DAP-Seq data were deposited in the NCBI Sequence Read Archive (Bioproject: PRJNA1271583) with the following BioSample: SAMN48877703 (L73-1087 Resequencing), SAMN48899944 (Lf2 DAP-Seq Rep1), SAMN48899945 (Lf2 DAP-Seq Rep2), SAMN48899946 (Input Control for DAP-Seq), SAMN48897410 (Clark Shoot Tip RNA Seq Rep 1), SAMN48897411 (Clark Shoot Tip RNA Seq Rep 2), SAMN48897412 (Clark Shoot Tip RNA Seq Rep 3), SAMN48897413 (L73-1087 Shoot Tip RNA Seq Rep 1), SAMN48897414 (L73-1087 Shoot Tip RNA Seq Rep 2), SAMN48897415 (L73-1087 Shoot Tip RNA Seq Rep 3).

## Acknowledgments

We would like to thank James Specht for providing us with the Clark *Lf2* near isogenic lines and Jinbin Wang for assistance in submitting samples for sequencing.

## Author Contributions

CBC and JM designed the research. CBC, DC, QZ, DP, ACE, QS, and CQ carried out the research. CBC, JM, DC, and ASIP analyzed the data. CBC wrote the manuscript with input from JM and the approval of all other authors.

## Supplemental Figure and Table Legends

**Supplemental Figure S1.** The first node after the cotyledon in L73-1087 (left) and Williams 82 (right) showing the normally simple unifoliate leaves becoming compound trifoliate leaves in the *lf2* mutant line

**Supplemental Figure S2.** Simple pedigree showing the relationship between L73-1087, T255, Clark, and Hawkeye.

**Supplemental Figure S3**. Protein Change in lf2 Mutant Lines. A Protein structure of wild-type Lf2 generated by Alpha-Fold. B Protein Structure of mutant lf2 generated by Alpha-Fold. The lack of the three helices in the bottom right corner corresponding to the homeobox DNA binding domain can be observed.

**Supplemental Figure S4**. T_1_ CRISPR-Cas9 Edited *lf2* Knockdown Lines Showing examples of four and five leaflets.

**Supplemental Figure S5**. RNA Sequencing Reads from Williams82 showing the annotated and alternative transcripts which allow the knockdown lines to maintain partial function.

**Supplemental Figure S6**. Top 10 overrepresented motifs from HOMER Analysis of Lf2 DAP-Seq Peaks

**Supplemental Figure S7**. DAP Seq Peaks in the *Lf1* Promoter

**Supplementary Table S1**. SNP Genotypes of Bulked Samples and Parents in the Region Defined By BSA

**Supplementary Table S2**. SNP Markers in the BSA Region from Publicly Available Resequencing Data Showing that the introgression of T255 DNA cannot be seen due to the close relationship among Clark and Hawkeye

**Supplementary Table S3**. Genes with Lf2 DAP-Seq Peaks in their Gene Bodies or Promoters

**Supplementary Table S4**. Differentially Expressed Genes Between the Clark *Lf2Lf2* and *lf2lf2* Near Isogenic Lines

