## Supplementary figures and images for "Lf2 is a knotted homeobox regulator that modulates leaflet number in soybean"

### FigS1

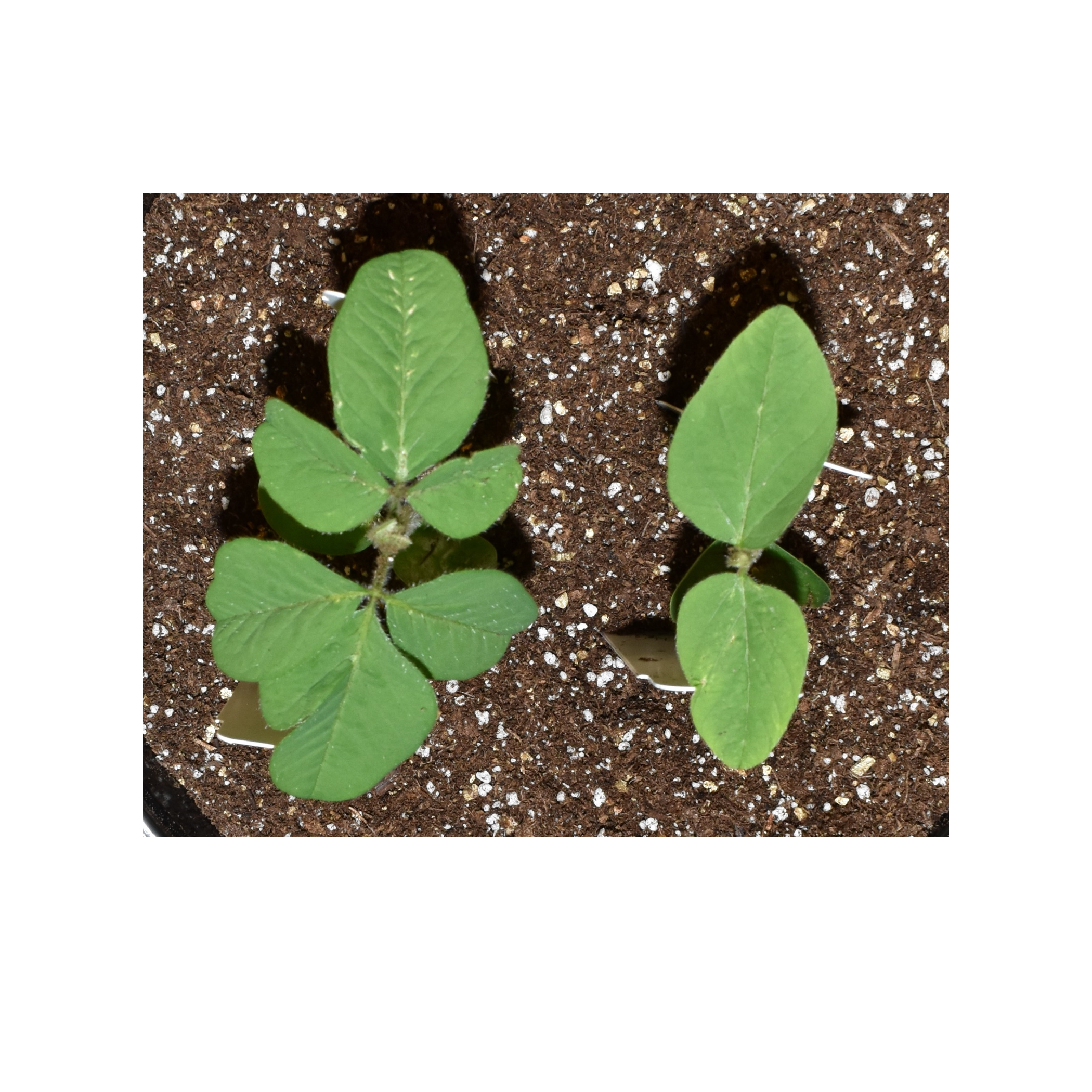

### FigS2

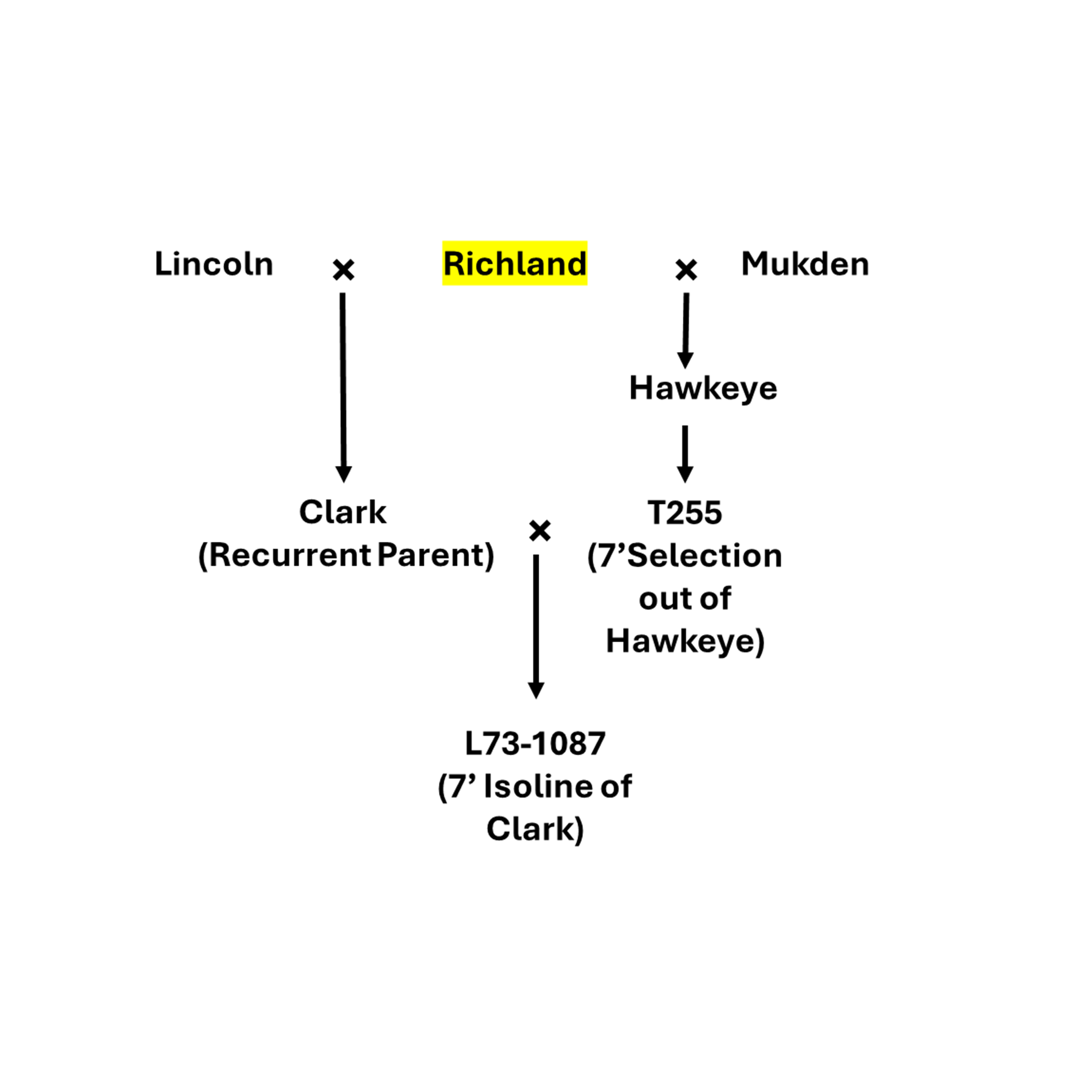

### FigS3

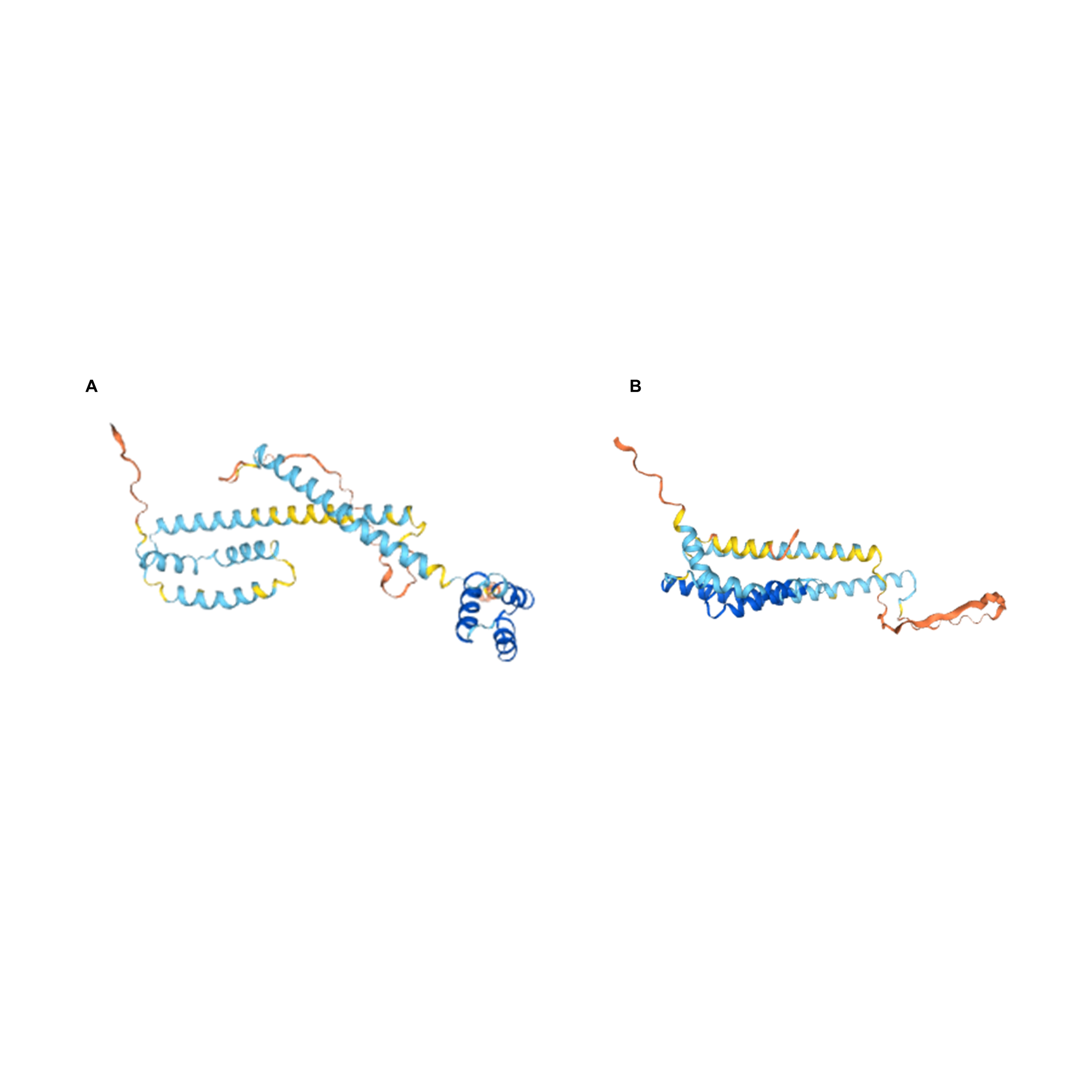

### FigS4

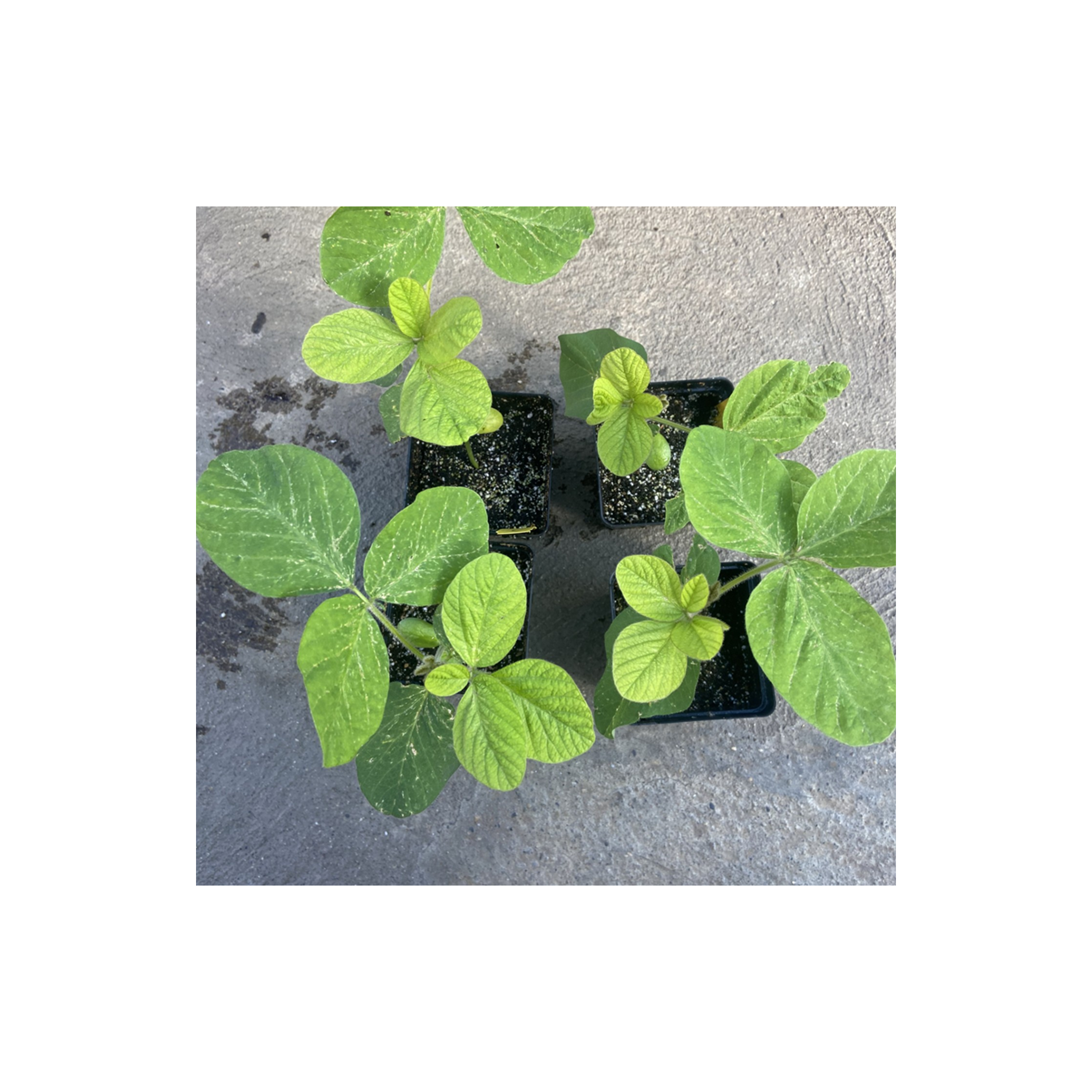

### FigS5

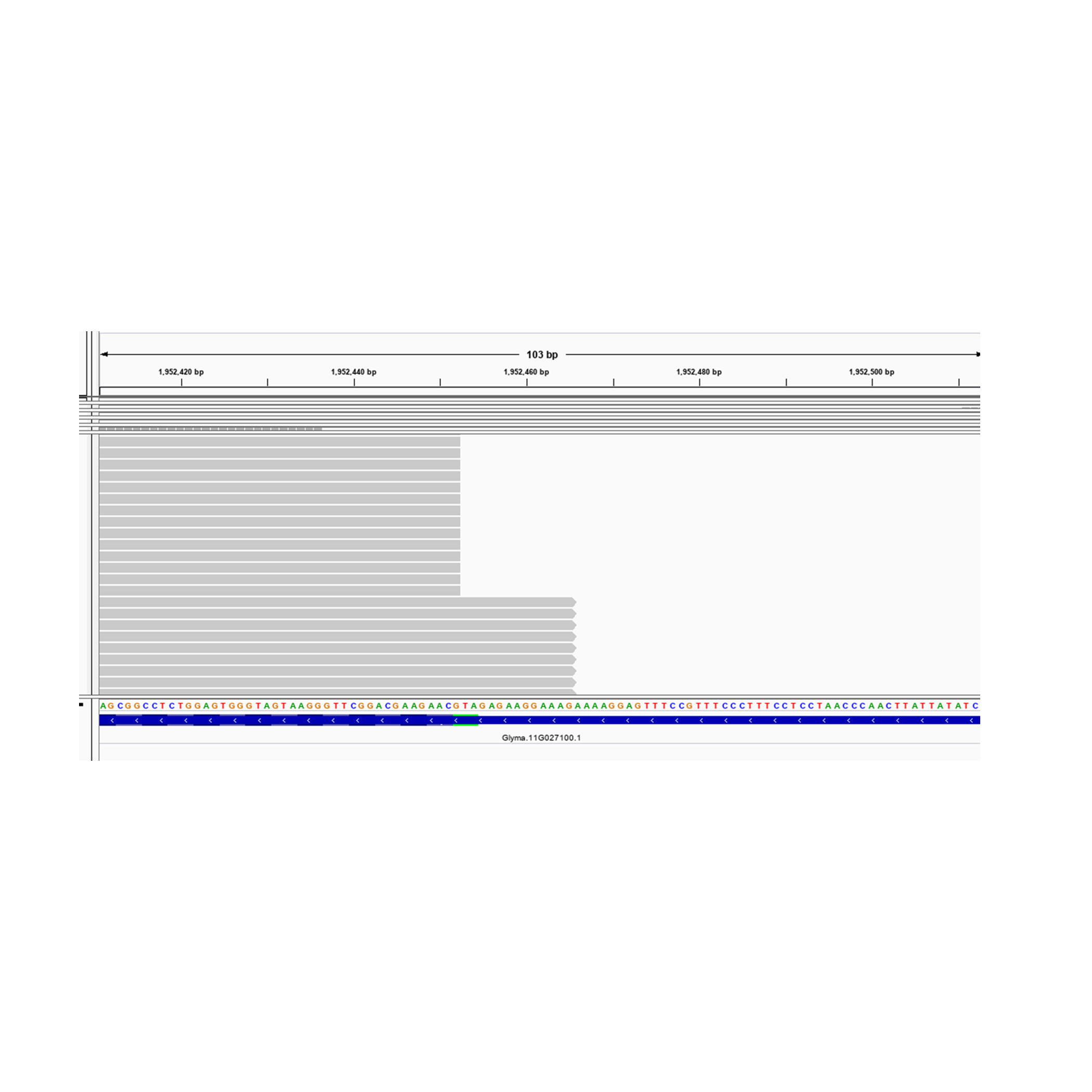

### FigS6

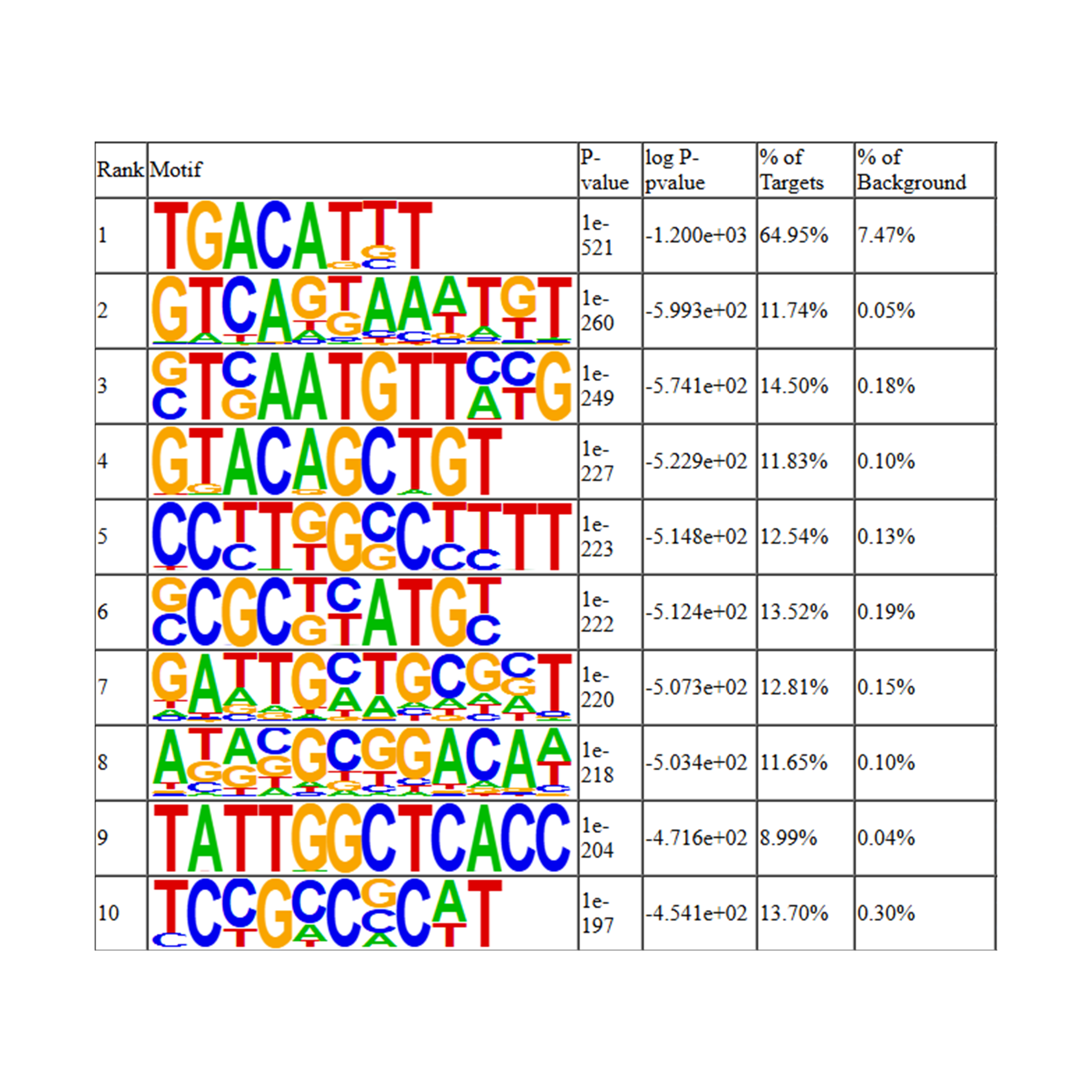

### FigS7

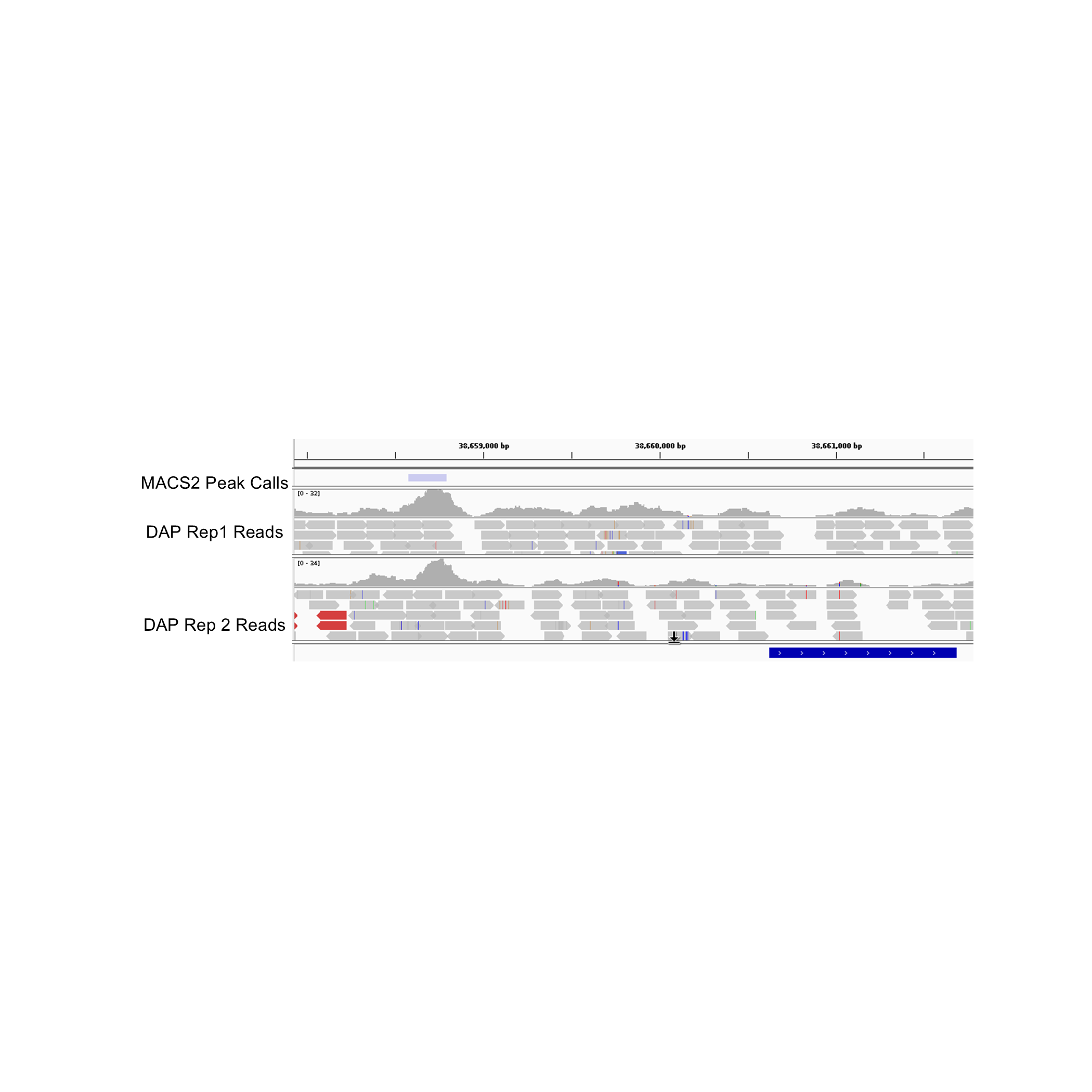
